# Bacterial and human exonucleases mediate interkingdom antiviral immunity

**DOI:** 10.64898/2026.06.30.735675

**Authors:** Courtney R. Santos, Alyaa El-Guindy, Aaron Embry, Arabella E. Martin, Neal M. Alto, Don B. Gammon, Kevin J. Forsberg

**Affiliations:** Department of Microbiology, University of Texas Southwestern Medical Center, Dallas, TX, USA

**Keywords:** Phage Defense, Innate Immunity, RNA Viruses, Evolution, Exonucleases, ISG20

## Abstract

All kingdoms of life have developed strategies to limit viral infection^1,2^. In humans, interferons induce a suite of antiviral factors that collectively provide immunity^3^. Some human immune genes are homologous with antiphage defense genes in bacteria, though the extent of this overlap is not known^4–12^. Here, we screened a panel of human innate immune genes for phage defense in *Escherichia coli* and found that the RNA exonuclease ISG20 potently restricts the RNA phages MS2 and Qβ. Purified ISG20 trims the 3’ untranslated regions (UTRs) of RNA phage genomes, explaining its ability to block phage replication in *E. coli*. Homologs of ISG20 from bacteria function similarly, exhibiting nuclease-dependent antiphage defense in bacteria and UTR-trimming *in vitro*. When expressed in human cells, these bacterial exonucleases also restrict human RNA viruses with similar potency as the human antiviral protein ISG20. Thus, antiviral genes from both humans and bacteria can function interchangeably. This bidirectional, interkingdom immunity suggests that viral targets overlap, implying that both bacterial and human factors recognize ancient features of RNA viruses.

## Introduction

The antiviral immune systems of humans were once considered completely distinct from those in bacteria. Recently, this dogma has been uprooted by the discovery of antiphage systems in bacteria homologous to antiviral defenses in humans^4–12^. These discoveries were made using DNA phages and suggest that similar antiviral strategies are conserved in bacteria and humans, despite substantial evolutionary divergence.

RNA viruses commonly infect mammals and bacteria, which may select for similar immune responses in each host^13–15^. To identify potential examples of this shared immunity, we expressed a subset of mammalian interferon-stimulated genes (ISGs) in *Escherichia coli* and screened for ones that confer defense against its phages (**Figure 1a**). We selected nine genes from *Homo sapiens* for functional testing, prioritizing genes with conserved interferon induction and demonstrated RNA interactions (**Extended Data Table 1**) ^16^. We intentionally omitted ISGs that require cofactors absent in *E. coli*, like ubiquitin, as we presumed these genes would be unlikely to function in our bacterial screening host.

**Fig. 1.**
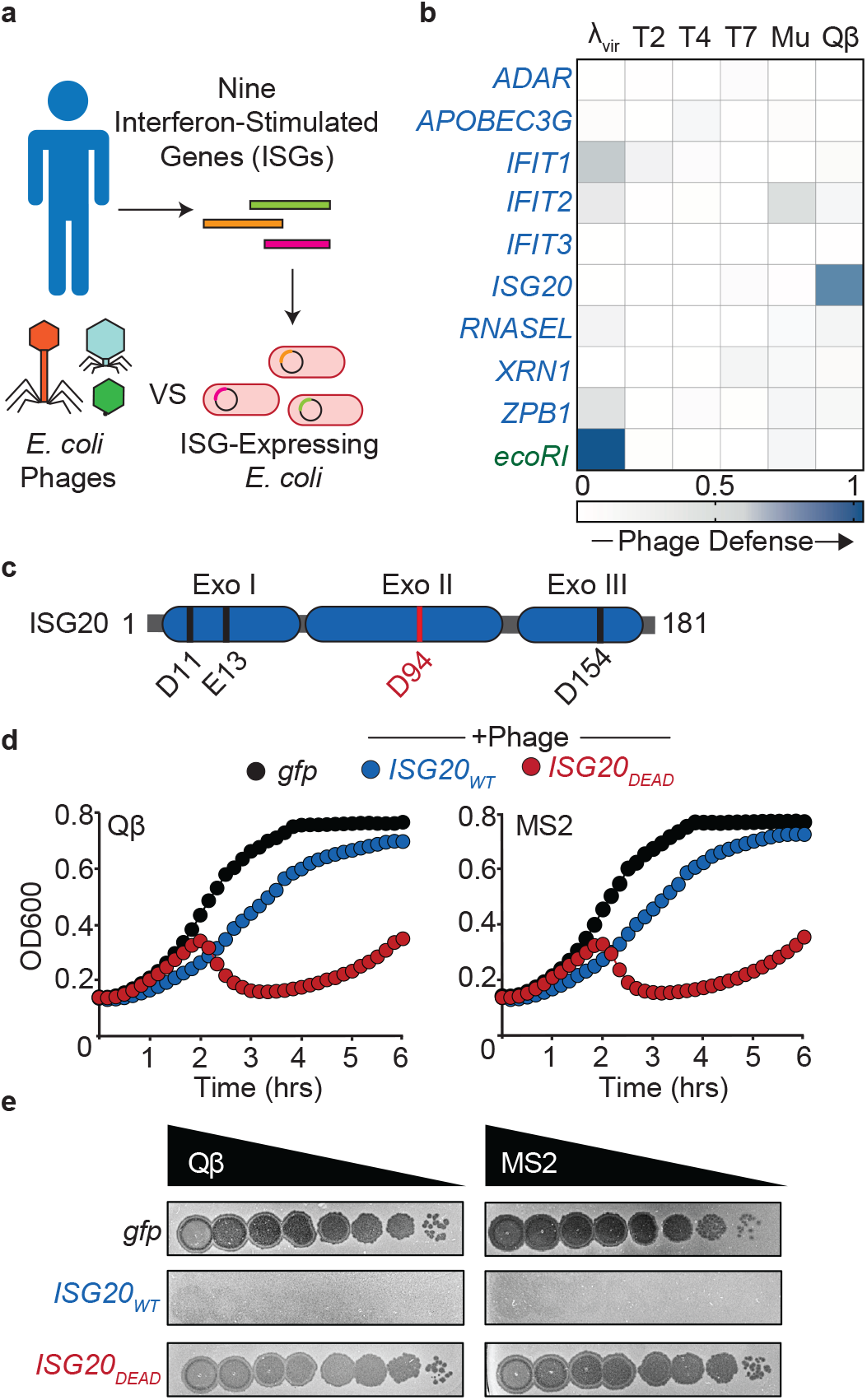
Human ISG20 protects *E. coli* from RNA phages. **a**, Schematic of a screen to identify mammalian ISGs that protect against phage infection. **b**, Heatmap depicting phage protection scores for ISGs (blue) and an *ecoRI* control (green) expressed in *E. coli*. **c**, ISG20 domain architecture with conserved catalytic residues mapped onto their respective exonuclease motifs (Exo). **d**, Growth curves of *E. coli* expressing *gfp, ISG20*_*WT*_, or *ISG20*_*DEAD*_ in the presence or absence of phage as indicated in the legend at top (MOI 0.1). Left, infection with phage Qβ; Right, infection with phage MS2. **e**, Double agar overlays depicting plaque phenotypes for Qβ (left) and MS2 (right) upon infecting *E. coli* expressing *gfp, ISG20*_*WT*_, or *ISG20*_*DEAD*_. Spots represent ten-fold dilutions from left to right. Growth curves depict averages of three technical replicates and are representative of three biological replicates. Error bars plot st. dev but are typically obscured by growth curve data points.

### Human *ISG20* prevents RNA phage infection when expressed in *E. coli*

To screen for antiphage defense, we exposed ISG-expressing *E. coli* to a panel of phages at various multiplicities of infection (MOI), monitoring bacterial growth over time via optical density. We compared each candidate gene to *gfp* controls with and without infection to approximate complete susceptibility and perfect defense, respectively. After phage infection, we assigned a ‘protection score’ to each ISG, calculated by normalizing bacterial growth to that of the uninfected *gfp* control (**Figure 1b, Extended Data Figure 1a,b,c**). As a positive control, we used the *ecoRI* restriction-modification system, which confers defense against phage λ^vir17^. This control produced a protection score near one whereas non-protective ISGs yielded scores closer to zero (**Figure 1b, Extended Data Figure 1b,c**). Of the nine ISGs tested, only one scored highly for phage defense. *ISG20*, which encodes a 3’ to 5’ RNA exonuclease, protected against Qβ, a phage with a positive-sense, single-stranded RNA (+ssRNA) genome (**Figure 1b,c and Extended Data Figure 1a**) ^18^.

We next tested whether *ISG20* conferred immunity against MS2, another +ssRNA phage that infects *E. coli*. In liquid growth curves, *ISG20* expression conferred phage defense against MS2 like it did for Qβ (**Figure 1d and Extended Data Figure 1a,d**). Cells expressing *ISG20* grew slightly slower than uninduced controls, indicating that this gene is only mildly toxic when expressed in *E. coli* (**Extended Data Figure 1e**). We also tested for phage defense via plaque enumeration on double agar overlays, finding that *ISG20* protected against both MS2 and Qβ, corroborating the results from our initial screen (**Figure 1e and Extended Data Figure 1f**).

### *ISG20* inhibits RNA phage replication

ISG20 is a DEDDh-family RNA exonuclease, named for conserved catalytic residues at the nuclease active site (**Figure 1c**). In mammals, ISG20 can defend against positive-sense and negative-sense RNA viruses, as well as some DNA viruses that replicate via genomic RNA intermediates^18–22^. Similarly, *ISG20* expression can protect *E. coli* from RNA phages, but not DNA phages (**Figure 1b,d,e**). Though ISG20’s mechanism of antiviral defense in mammals is not certain, and may vary with cell type and virus, the catalytic residue Asp94 is typically required for immune function^18–22^. We also found that expression of a D94A mutant (hereafter *ISG20*^*DEAD*^) is incapable of defending *E. coli* from the phages MS2 and Qβ (**Figure 1d,1e and Extended Data Figure 1f**). Thus, as is the case in mammals, ISG20’s nuclease activity is required for defense against RNA viruses in bacteria.

To infect *E. coli*, both MS2 and Qβ adsorb to the F pilus, a critical component of the Type IV secretion system encoded by the conjugative F plasmid in *E. coli* (**Figure 2a**). As a result, these phages can only infect F(+) *E. coli* strains, which develop resistance to infection if this pilus is downregulated or inactivated^23^. To determine if *ISG20* interferes with receptor attachment, we measured MS2 and Qβ adsorption to an F(+) strain in the presence and absence of *ISG20*. As controls, we included an F(−) strain and an F(+) strain expressing *finO* instead of *ISG20*, as *finO* represses the F pilus and prevents phage adsorption^24,25^. As expected, both phages adsorbed to F(+) cells in the absence of *ISG20*, co-pelleting with these cells after centrifugation and reducing phage titers in culture supernatants (**Figure 2b**). Non-adsorptive controls also matched expectation, as neither F(−) nor F(+)-*finO* cells co-pelleted with RNA phages. Notably, phage adsorption in *ISG20*-expressing cells was indistinguishable from wild-type F(+) controls. We therefore conclude that *ISG20* acts downstream of adsorption to block MS2 and Qβ infection (**Figure 2b**).

**Fig. 2.**
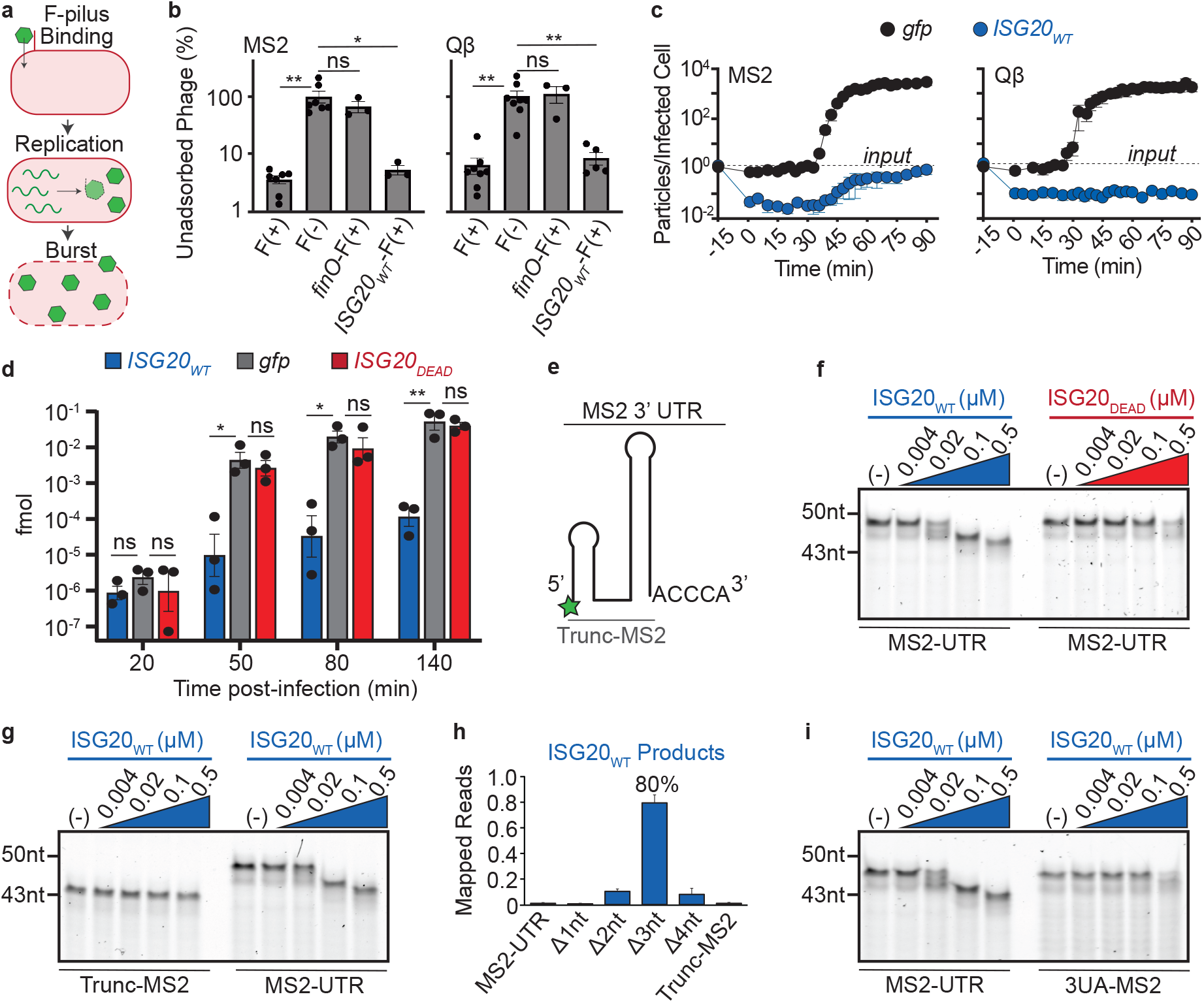
ISG20 blocks MS2 replication and degrades MS2 3’ UTRs. **a**, Schematic depicting the MS2 infection cycle. **b**, Phage recovered from culture supernatants after adsorption to *E. coli* expressing *finO, ISG20*_*WT*_, or neither at MOI 1. The F(−) strain does not encode the F-pilus and is the comparator for tests of statistical significance. *p≤0.05; **p≤0.005, one-way ANOVA and Dunnett’s post-test. Error bars depict standard error of the mean. **c**, Enumeration of infectious MS2 (left) or Qβ (right) phage particles, produced over time from cells expressing either *gfp* or *ISG20*_*WT*_. Phages were allowed to adsorb to the indicated host for 15 minutes before sampling. **d**, Quantification of MS2 genomic RNA levels over time by qRT-PCR, from infections of *E. coli* cells expressing *gfp, ISG20*_*WT*_, or *ISG20*_*DEAD*_. *p≤0.05; **p≤0.005, two-way ANOVA and Dunnett’s post-test. **e**, Diagram of the terminal region of MS2’s 3’ UTR, synthesized with a 5’ 6-FAM fluorophore (green star). A truncated form without the final five ribonucleotides was also synthesized, as indicated. **f**, Degradation of the MS2-R UTR (0.5µM) after 15-minute incubations with either ISG20_WT_ or ISG20_DEAD_, at the indicated protein concentrations. **g**, Degradation products from the Trunc-MS2 and MS2-UTR substrates (0.5µM) incubated with ISG20_WT_, as in panel **f. h**, Sequencing reads of MS2-UTR degradation products after incubation with ISG20_WT_. **i**, Degradation products from the MS2-UTR and 3UA-MS2 substrates (0.5µM) incubated with ISG20_WT_, as in panel **f**. RNA gels depicted in panels **f, g**, and **i** are representative of three biological replicates. Error bars depict standard error of the mean for 3 biological replicates.

We next performed one-step phage infections to measure MS2 and Qβ burst size in the presence of *gfp*- and *ISG20*-expressing *E. coli* cells. In line with previous reports, both phages produced about 10^3^ infectious particles for every input phage after one round of infection on the permissive host^26^. In contrast, when mixed with *ISG20*-expressing cells, both MS2 and Qβ yielded fewer infectious particles than input (**Figure 2c**). Therefore, *ISG20* inactivates these +ssRNA phages. To determine if expressing the enzyme prevents genome replication, we measured MS2 genomic RNA levels via qRT-PCR (**Figure 2d**). In cells expressing *gfp* or *ISG20*_*DEAD*_, we observed a >1000-fold increase in genomic RNA during MS2 infection. In comparison, little to no replication of MS2 genomic RNA was observed in *ISG20*-expressing cells. Thus, human *ISG20* blocks MS2 replication in *E. coli* and requires an intact nuclease domain to mediate its antiviral effect.

To evaluate *ISG20*’s mechanism, we first asked whether it elicited phage defense via abortive infection. In abortive infection defense systems, an infected cell initiates programmed cell death or dormancy to prevent phage replication and protect neighboring cells^27^. At low MOI, population growth is unaffected, as infected cells are rare. However, at high MOI, every cell is infected and initiates programmed cell death or dormancy, with drastic impacts on total population growth. To test whether *ISG20* induces this response upon phage infection, we subjected *ISG20*-expressing *E. coli* to MS2 and Qβ infection at high (10) and low (0.1) MOI. Cells expressing *ISG20* maintain antiphage activity and exhibit similar growth curves at both MOIs, indicating that phage defense does not occur via abortive infection (**Extended Data Figure 1g,h**). We therefore evaluated whether ISG20 elicits immunity via direct action against the infecting phage.

### ISG20 degrades RNA phage 3’ UTRs

Because ISG20 is a 3’ to 5’ RNA exonuclease, we considered phage RNAs with exposed 3’ ends as potential immune targets. Both MS2 and Qβ have highly conserved 3’ UTRs, which are critical for initiating replication, packaging genomes into phage capsids, and other essential processes^28–30^. For instance, the 3’-CCCA end of Qβ’s genome is essential for initiation of RNA synthesis. Loss of just the terminal adenosine reduces RNA polymerization tenfold, and loss of the final two bases prevents phage replication^31–33^. To examine potential interactions with ISG20, we synthesized the terminal ends of each phage’s 3’ UTR with a 5’ fluorophore label. In our designs, we preserved each UTR’s final two stem-loops and their - CCCA 3’ ends to approximate native RNA secondary structures (**Figure 2e and Extended Figure 2a**). We incubated these RNAs with purified ISG20 and noticed a rapid degradation of their 3’ ends with sub-stoichiometric protein levels (**Figures 2f and Extended Data Figure 2b**). This degradation was not observed with purified ISG20_DEAD_, indicating that nuclease activity is attributable to ISG20 (**Figures 2f and Extended Data Figure 2b**).

Notably, ISG20’s degradation product was only a few bases smaller than the initial substrate, implying that the enzyme could not processively degrade the entire UTR. This result is consistent with ISG20’s distributive nuclease activity and inability to degrade structured RNA^34,35^. To probe further, we synthesized a truncated version of the MS2 3’-UTR (Trunc-MS2) and a variant that swaps the stem sequences of the terminal loop, altering UTR sequence while preserving structure (stem-swap, or SS-MS2). We observed that Trunc-MS2 was resistant to ISG20 degradation, indicating that ISG20 acts only on its single-stranded, 3’-CCCA overhang (**Figure 2g,h**). In contrast, the SS-MS2 substrate was degraded at rates comparable to the wild-type MS2 sequence, suggesting that ISG20 activity is not dependent on stem sequence (**Extended Data Figure 2c**). To test for sequence dependence on the 3’-CCCA overhang, we changed it to 3’-UUUA (3UA-MS2) and repeated the nuclease assay. The 3UA-MS2 substrate resisted ISG20 degradation compared to the wild-type MS2 3’ UTR, indicating that ISG20 has some sequence preference for MS2’s terminal cytosines (**Figure 2i**).

To determine the exact sequence of ISG20’s degradation product, we sequenced MS2’s wild-type UTR following incubation with ISG20. In over 99% of sequencing reads, at least one ribonucleotide was truncated from the 3’ end of the RNA, consistent with complete elimination of the full-length substrate (**Figure 2h**). In 80% of truncated RNAs, exactly three ribonucleotides were removed, indicating that this is the major product of ISG20’s exonuclease activity (**Figure 2h**). These ribonucleotides are strictly required for replication of RNA phage genomes, and so can explain ISG20’s antiphage activity in bacteria^30–33^. Consistent with this model, ISG20 severely suppresses replication of MS2 genomic RNA *in vivo*, in a nuclease-dependent manner (**Figure 2d**).

### *ISG20*-like homologs from bacteria defend against RNA phages

Given ISG20’s activity against RNA phages, we wondered whether related DEDDh-family exonucleases from bacteria could exert a similar effect. To identify bacterial homologs of ISG20 for functional testing, we searched NCBI’s nr, nt, and conserved domain databases, using human ISG20 as a query. We selected 27 sequences for functional interrogation that spanned the phylogenetic distribution of these hit lists, with some preference for sequences in γ-proteobacteria (**Figure 3a, Extended Data Table 2**). As expected, structural predictions of these homologs most closely resembled DEDDh-family exonucleases, including examples that degrade DNA and RNA substrates (**Extended Data Table 3**). On a sequence level, these genes were clearly distinct from other DEDDh exonuclease domains associated with phage defense^36,37^. They also co-localized with other predicted defense genes and within mobile genetic elements like prophages, suggesting a native defense function (**Figure 3b**). One example was encoded by a hybrid IncF/IncC plasmid from *E. coli*, which is closely related to the F plasmids that confer susceptibility to RNA phages^38^. Given these associations with phage defense, we tested whether bacterial homologs of ISG20 could defend against RNA phages in *E. coli*.

**Fig. 3.**
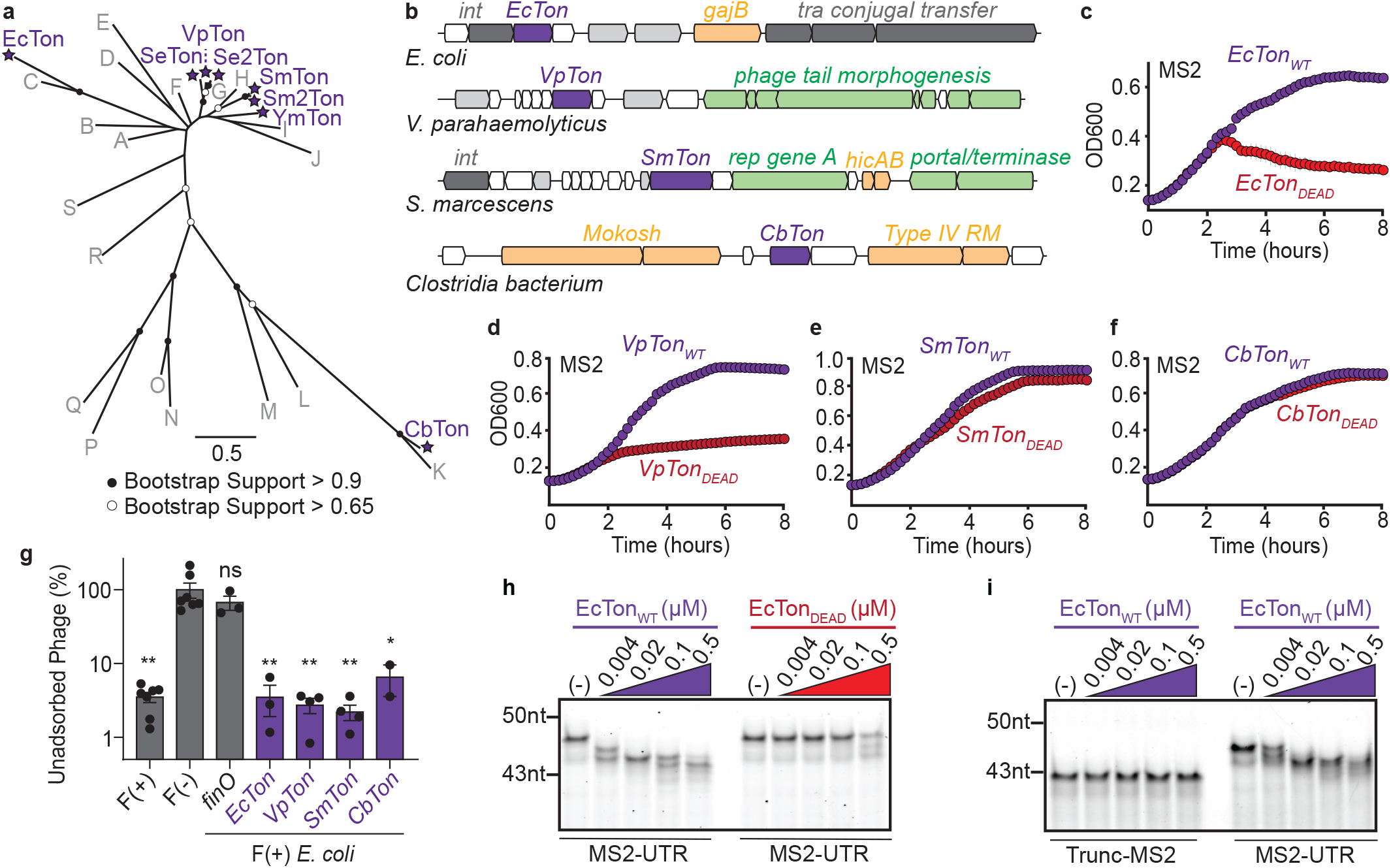
Bacterial homologs of ISG20 protect *E. coli* from MS2 infection. **a**, Unrooted phylogenetic tree depicting 27 putative ISG20 homologs from bacterial genomes. Circles at nodes indicate bootstrap support >0.9 or >0.65 if filled or empty, respectively. Scale bar indicates the number of amino acid substitutions per site. Sequences that defend against MS2 are named and labeled with purple stars; gray letters denote non-protective sequences. **b**, Genomic context for selected TONDO genes, colored in purple. Nearby genes are colored by function: yellow indicates phage defense, green indicates other phage functions, dark gray indicates integrases or conjugative pilus machinery, light gray indicates genes with other known functions, and white indicates genes with unknown functions. **c-f**, Growth curves of *E. coli* expressing the indicated TONDO (purple) or its catalytically inactive mutant (red), in the presence of MS2 at an initial MOI of 0.1. **c**, MS2 defense by *EcTon*_*WT*_ and *EcTon*_*DEAD*_. **d**, MS2 defense by *VpTon*_*WT*_ and *VpTon*_*DEAD*_. **e**, MS2 defense by *SmTon*_*WT*_ and *SmTon*_*DEAD*_. **f**, MS2 defense by *CbTon*_*WT*_ and *CbTon*_*DEAD*_. **g**, MS2 recovered from culture supernatants after adsorption to *E. coli* expressing *finO* or the indicated TONDO gene (purple), at MOI 1. The F(−) strain does not encode the F-pilus and is the comparator for tests of statistical significance. *p≤0.05; **p≤0.005, one-way ANOVA and Dunnett’s post-test. Error bars depict standard error of the mean. **h**, Degradation of the MS2-UTR (0.5µM) after 15-minute incubations with either EcTon_WT_ or EcTon_DEAD_, at the indicated protein concentrations. **i**, Degradation products from the Trunc-MS2 and MS2-UTR substrates (0.5µM) incubated with EcTon_WT_, as in panel **h**. Growth curves depict averages of three technical replicates. Error bars plot st. dev. but are typically obscured by growth curve data points. Growth curves and RNA gels are representative of three biological replicates.

To identify homologs with antiphage activity, we expressed genes encoding each candidate in *E. coli* and challenged these bacteria with MS2. Of the 27 predicted exonucleases tested, eight protected against infection (**Figure 3c,d,e,f and Extended Data Figure 3**). We named these ISG20 homologs TONDOs (<u>T</u>erminal <u>O</u>ligoribo-<u>N</u>ucleases with <u>D</u>efense <u>O</u>utcomes), based on their antiphage activity, enzymatic function, and in homage to the populous district in Manila, Philippines. As with *ISG20*, TONDOs did not impact phage adsorption and did not exhibit MOI-dependent phenotypes suggestive of abortive infection (**Figure 3g and Extended Data Figure 4a,b,c,d**). To determine if exonuclease activity was required for phage defense, we made catalytic mutations in four examples (D191A in *EcTon*_*DEAD*_, D100A in *VpTon*_*DEAD*_, D100A in *SmTon*_*DEAD*_, and D91A in *CbTon*_*DEAD*_). In two cases (*EcTon*_*DEAD*_ and *VpTon*_*DEAD*_), cells expressing active site mutants became susceptible to MS2 and Qβ infection (**Figure 3c,d and Extended Data Figure 4e,f**). In two others (*SmTon*_*DEAD*_ and *CbTon*_*DEAD*_), cells expressing these mutants remained completely or partially resistant to phage infection (**Figure 3e,f and Extended Data Figure 4g,h**). Thus, nuclease activity is sometimes dispensable for phage defense among TONDOs. This prompted us to examine other features of these enzymes.

Unlike ISG20, which contains only a standalone DEDDh exonuclease domain, TONDOs typically contain an auxiliary domain, which differs by homolog and often lacks recognizable function (**Extended Data Figure 4i**). Because ISG20 partners with different auxiliary proteins to mediate immunity against different infecting viruses^21^, we wondered whether these auxiliary domains may similarly potentiate antiphage defense. We noticed that one putative exonuclease conspicuously failed to defend against RNA phage (Candidate G), despite high sequence similarly to three functional TONDOs (SeTon, Se2Ton, VpTon; **Figure 3a and Extended Figure 4j**). Unlike these functional TONDOs, Candidate G lacks an auxiliary domain. However, this gene is immediately adjacent to another small open reading frame (ORF) in its native genome context (**Extended Data Figure 4k**). When co-expressed with this auxiliary ORF, candidate G conferred robust defense against MS2 and so was renamed *VΦTon* (**Extended Data Figure 4l**). Neither *VΦTon* nor the auxiliary ORF was functional in isolation (**Extended Data Figure 4m,n**). Evidently, like many phage defense systems, TONDOs use modular domains and genes to exert phage defense.

**Fig. 4.**
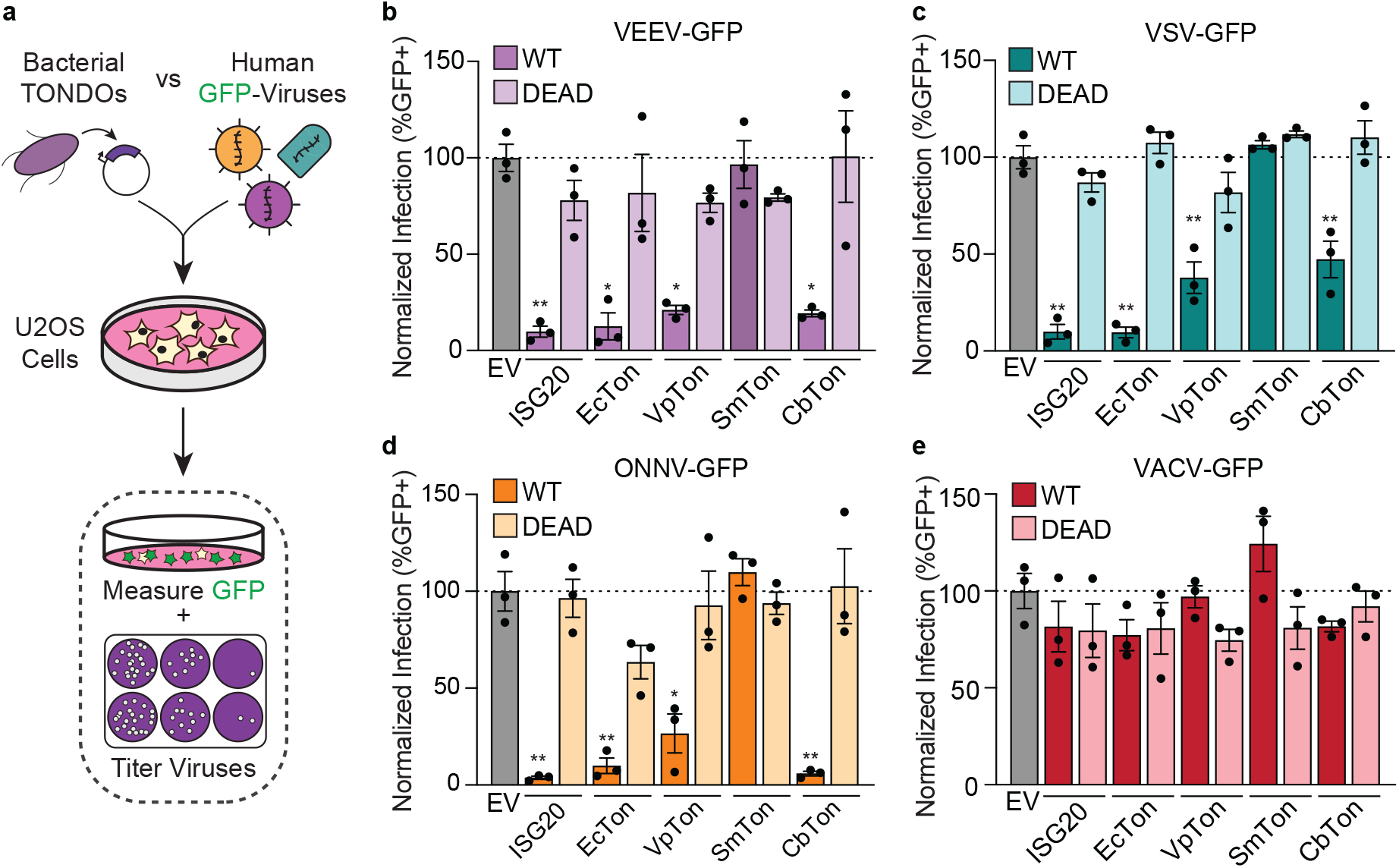
TONDOs confer antiviral immunity in human cells. **a**, Experimental scheme used to test for antiviral immunity. Successful infections will yield GFP+ cells and new viral progeny. **b**, Viral GFP signal relative to empty vector (EV) control cells 24 hours post-infection (hpi) with VEEV-GFP (MOI 1). **c**, Viral GFP signal relative to EV control cells 24 hpi with VSV-GFP (MOI 1). **d**, Viral GFP signal relative to EV control cells 24 hpi with ONNV-GFP (MOI 0.5). **e**, Viral GFP signal relative to EV control cells 24 hpi with VACV-GFP (MOI 1). The data represent means of three biological replicates; error bars depict st. dev. Statistical significance determined by one-way ANOVA and Dunnett’s post-test, compared to EV controls; *p≤0.05, **p≤0.005. Comparisons without asterisks are not statistically significant. Viral titers are shown in **Extended Data Figure 6**.

We next investigated whether TONDOs act similarly to ISG20 *in vitro*. We chose to interrogate *EcTon* based on its strong phage defense, presence on IncF-like plasmids, and because, like *ISG20*, its gene exhibited nuclease-dependent activity *in vivo* (**Figure 3c**). We purified EcTon and EcTon_DEAD_, incubating these proteins with MS2 and Qβ 3’ UTRs to test for potential activity. As we observed for ISG20, EcTon rapidly degraded the 3’ end of MS2 and Qβ UTRs at sub-stoichiometric concentrations (**Figure 3h and Extended Data Figure 5a**). Little to no activity was observed for EcTon_DEAD_, consistent with *EcTon*’s nuclease-dependent activity *in vivo* (**Figure 3h and Extended Data Figure 5a**). Like ISG20, EcTon was inhibited by structured RNA ends, as it could not degrade Trunc-MS2 (**Figure 3i**). Similarly, EcTon degraded SS-MS2 at comparable rates to the WT-MS2 UTR, indicating that its activity is not dependent on UTR stem sequence (**Extended Data Figure 5b**). Unlike ISG20, EcTon showed no preference for MS2’s native 3’-CCCA overhang, as it degraded the 3UA-MS2 substrate as efficiently as the wild-type UTR (**Extended Data Figure 5c**).

### Bacterial TONDOs defend against human RNA viruses

Given that ISG20, a human antiviral protein, could provide phage defense when expressed in bacteria, we wondered whether the reverse was true for bacterial TONDOs. We codon-optimized four TONDOs for expression in mammalian cells along with their corresponding nuclease-dead mutants. We then ectopically expressed these eight genes in human U2OS osteosarcoma cells, comparing their phenotypes to those of *ISG20*_*WT*_, *ISG20*_*DEAD*_, and empty-vector controls. The expression of each protein was detected, with DEAD mutants at higher levels than their wild-type counterparts, consistent with reported literature (**Extended Data Figure 5d**) ^39^. We then tested each construct for antiviral defense against four GFP-tagged mammalian viruses: +ssRNA Venezuelan Equine Encephalitis Virus (VEEV), +ssRNA Onyong-nyong Virus (ONNV), −ssRNA Vesicular Stomatitis Virus (VSV), and Vaccinia virus (VACV), which has a DNA genome^40–42^. This set includes RNA viruses known to be restricted by ISG20 (VEEV and VSV), a DNA virus for which restriction is not anticipated (VACV), and an RNA virus for which ISG20 susceptibility is unknown (ONNV) ^18–20^. After expressing putative antiviral factors for 24 hours, cells were infected for an additional 24 hours and viral replication was quantified by both GFP expression and plaque assay to determine titer (**Figure 4a**). For the duration of the experiment, there were no cell viability differences among the various constructs (**Extended Data Figure 5e**).

As expected, expression of *ISG20*_*WT*_ inhibited infection by the RNA viruses VEEV and VSV (**Figure 4b,c and Extended Data Figure 6**). The +ssRNA virus ONNV was also restricted by *ISG20*_*WT*_ expression, suggesting that it is susceptible to this antiviral effector like its related alphavirus, VEEV (**Figure 4d and Extended Data Figure 6**). In contrast, *ISG20*_*DEAD*_ was unable to restrict any of the RNA viruses, and VACV was unaffected by either *ISG20*_*WT*_ or *ISG20*_*DEAD*_ (**Figure 4b,c,d,e and Extended Data Figure 6**). Similarly, neither *SmTon*_*WT*_ nor *SmTon*_*DEAD*_ could restrict any virus in the panel. Remarkably, ectopic expression of *EcTon*_*WT*_, *VpTon*_*WT*_, and *CbTon*_*WT*_ each restricted all RNA viruses tested, with their nuclease-dead mutants exhibiting no antiviral phenotypes (**Figure 4b,c,d,e and Extended Data Figure 6**). Thus, these TONDOs restrict human RNA viruses in a nuclease-dependent manner, like *ISG20*. Both *VpTon*_*WT*_ and *CbTon*_*WT*_ restricted the +ssRNA viruses VEEV and ONNV more potently than the −ssRNA virus, VSV, perhaps related to their activity against +ssRNA phages in bacteria. Strikingly, expression of the *E. coli* protein EcTon_WT_ protected mammalian cells from both −ssRNA and +ssRNA viruses as potently as the human antiviral effector, ISG20 (**Figure 4b,c,d,e and Extended Data Figure 6**). These results reveal that antiphage exonucleases from bacteria can provide broad-spectrum antiviral immunity when expressed in human cells.

## Discussion

By screening human antiviral genes in *E. coli*, we learned that *ISG20* potently defends bacteria against RNA phages. ISG20 is a 3’ to 5’ RNA exonuclease and requires nuclease activity to mediate phage defense in *E. coli*, like it does for antiviral immunity in mammals. This nuclease activity is also needed to trim phage 3’ UTRs *in vitro*, which can explain ISG20’s ability to block phage genome replication *in vivo*. Homologs of ISG20 from bacteria function similarly, with examples that trim UTRs and block phage infection in a nuclease-dependent manner. Their genes reside in prophages and other mobile genetic elements, which suggests a native role in phage defense. Like their human counterpart, they also exhibit interkingdom antiviral immunity. Three of these bacterial exonucleases restrict RNA viruses when expressed in human cells, including one example from *E. coli* that blocked infection as potently as *ISG20*.

In humans, ISG20’s exact mechanisms of antiviral defense are unclear. It confers immunity against diverse RNA viruses via partnerships with different condition- or virus-specific cofactors. This flexibility enables different defense strategies to be tailored to specific viral threats and infection contexts^18^. In some cases, ISG20 acts directly on viral RNAs, including their 3’ UTRs^21,22^. In others, ISG20 blocks infection by indirect action, for instance by impacting translation or stimulating expression of other antiviral factors^20,43^. Bacterial homologs of ISG20 may be similarly diverse, enacting different defense strategies via their often-unrelated auxiliary domains. Presumably, these mechanisms sometimes target viral components or host processes that are conserved across kingdoms, enabling their action against RNA viruses in human cells.

Two groups have shown that human ISGs can defend bacteria from DNA phages, by targeting viral features conserved across kingdoms^6,44^. We show that the same phenomenon occurs with RNA phages, for which few antiphage defenses are known^15^. We also demonstrate that interkingdom antiviral immunity can be bidirectional: immune genes from humans and bacteria each function in both hosts interchangeably. Though host-virus competition stimulates rapid change, it also replays the same themes and reuses the same components across evolutionary timescales^45,46^. In the case of ISG20, it seems that both humans and bacteria are reading from the same page of the antiviral playbook.

## Methods

### Expression constructs and bacterial handling

Human ISGs were codon-optimized for *E. coli*, synthesized by GenScript, and cloned into the pZE21_TetR vector for inducible expression. Bacterial *ISG20* homologs were synthesized and cloned identically; all sequences are listed in **Extended Data Table 5**. Before phage defense or bacterial growth assays, plasmids were chemically transformed into NEB Turbo *E. coli* cells and single colony transformants were grown overnight in LB broth (10 g/L casein peptone, 10 g/L NaCl, 5 g/L ultra-filtered yeast powder) supplemented with kanamycin (50 μg mL^−1^), IPTG (0.5 mM), MgCl_2_(1.25 mM), and CaCl_2_(1.25 mM). The next day, 2% outgrowth cultures were made in 4-5 mL using the same medium as was used for overnight cultures, additionally supplemented with doxycycline (100 ng µL^-1^) to induce gene expression. Cultures were grown at 37 °C with shaking at 200rpm until mid-logarithmic growth was achieved (OD_600_ values 0.3 – 0.7), unless otherwise stated.

### Phage infections

The phages used in this study were described previously^46^. Phage details are as follows: T4 (courtesy of Dr. Frederick Bushman, NCBI KJ477684.1), T2 (ATCC 11303-B2), T7 (ATCC BAA-1025-B2), λ_vir_ (courtesy of Dr. Gerry Smith), Qβ (ATCC 23631-B1), and MS2 (ATCC 15597-B1).

Liquid media infections and bacterial growth curve assays were performed using GFP, Human ISGs, bacterial TONDOs, candidate genes, control genes, and nuclease-dead mutants of these constructs. Mid-log cultures were diluted to OD_600_ = 0.2 in either inducing or non-inducing conditions, as described, with subsequent phage addition (at listed MOIs). Samples were split into three technical replicates of 100 µL and dispensed in technical triplicate into flat-bottom 96-well plates (Costar). OD_600_ was measured over time on the LogPhase 600 (Agilent Bio Tek) with the following settings: [Discontinuous Kinetics: Runtime 01:00:00 (D:HH:MM), Interval 00:10:00 (HH:MM:SS), 145 Reads (max) Read: 600 nm Incubator: On Temperature setpoint: 37 °C Temperature gradient: On Shaking: On Shaking frequency: 800 RPMs]. For the initial ISG screen, phage protection scores were calculated relative to the OD_600_ values of infected and uninfected *gfp* controls cells (scored as 0 and 1, respectively). For each phage listed in **Figure 1b**, OD_600_ values were used at consistent timepoints and MOIs to calculate these scores. Timepoints and MOI pairs are as follows for each phage: [λ_vir_ (6hrs, MOI 10^−3^); T4 (4hrs, MOI 10^−5^); T7 (9hrs, MOI 10^−4^); T2 (8hrs, MOI 10^−4^); Qβ (5hrs, MOI 10^−1^); Mu (5.5hrs, MOI 10^−1^)].

MOIs were chosen based on the lowest value that consistently triggered culture crash for each phage with *gfp* controls, with timepoints selected at the trough of OD_600_ values for these growth curves. For other liquid phage infections and growth assays, curves reported are representative of three independent biological replicates unless noted.

Double agar plaque assays were performed using *gfp, ISG20*_*WT*_, and *ISG20*_*DEAD*_ expressed as described above. Outgrowth cultures were grown to an OD_600_ >1.0 (typically 1.1-1.6). Then, 200 µL of cells were mixed with 3 mL of LB top-agar [LB Broth with 0.5% agar pre-supplemented with MgCl_2_(1.25 mM) and CaCl_2_(1.25 mM), doxycycline (100 ng uL^-1^)], and then immediately poured onto 1% LB-agar plates containing kanamycin (50 μg mL^−1^), IPTG (0.5 mM), and doxycycline (100 ng uL^-1^) before drying top agar by flame for about 5 minutes. Tenfold serial dilutions of Qβ or MS2 phage were then spotted in technical triplicate across each plate and dried by the flame for another 5 minutes before incubating at 37 °C overnight. Phage protection was quantified by counting plaque forming units (PFUs) and calculating efficiency of plaquing (EOP) relative to *gfp* controls.

### Adsorption Assays

Constructs containing *gfp, finO, ISG20*_*WT*_, or bacterial TONDOs were expressed and passaged as described in the first paragraph of the Methods section. As an F(−) strain control, *gfp* constructs were also transformed and expressed in *E. coli* MegaX (ThermoFisher cat. #C640003). Mid-log cells were harvested by centrifugation (4121g, 10 min), resuspended in fresh LB broth supplemented with MgCl_2_(1.25 mM) and CaCl_2_(1.25 mM), and infected with phage at an MOI of 1. Samples were incubated at 37 °C with shaking for 15 min to allow phages to adsorb to cells. Bacterial cells, along with adsorbed phages, were pelleted (4000g, 10 min) and the supernatants containing unadsorbed phages were filtered through 0.22-μm syringe filters to remove residual bacterial debris. Phage titers of the filtered supernatants were determined by double-agar plaque assays using NEB Turbo as an indicator strain, as described above. The percentage of unadsorbed phage PFUs compared to input is plotted.

### RT-qPCR

Constructs containing *gfp, ISG20*_*WT*_, and *ISG20*_*DEAD*_ were expressed and passaged as described in the first paragraph of the Methods section. Cultures were normalized to an OD_600_ of exactly 0.4 and incubated at 37 °C for 5 minutes. Then, MS2 was added at an MOI of 0.1 and incubated at 37 °C, without shaking. Equal volumes of SM buffer (100mM NaCl, 50 mM Tris-Cl, 2 mg/mL MgSO_4_·7H_2_O) was added to uninfected control samples. At given timepoints, infected or uninfected cells were added to chilled 10X Stop Buffer (95% ethanol, 5% TRIzol) and immediately inverted to mix and stop infection. Samples were centrifuged (13000rpm, 30 sec) and supernatants discarded. Pellets were resuspended in 1X Stop Buffer (diluted in 10x PBS) before centrifugation (13000rpm, 30 sec). Then, supernatants were discarded, and pellets were subsequently flash frozen in liquid nitrogen until sample collection was completed. To extract RNA, 400 µL of 65 °C TRIzol was added to frozen pellets, vortexed, and incubated in a thermomixer set at 65 °C for 10 minutes, shaking at 2000 rpm. The samples were then flash frozen in liquid nitrogen for at least 10 minutes, thawed at room temperature, and then centrifuged in a tabletop centrifuge (21300g, 5 min) at 4 °C. The TRIzol-treated supernatants were added to 400 µL of 100% ethanol and processed using the Direct-zol RNA miniprep kit (Zymo Research cat. #R2050) following manufacturer’s instruction. Briefly, samples were applied to spin columns and centrifuged before transferring columns to fresh collections tubes. Columns were treated with PreWash and RNA Wash Buffer before eluting RNA in 88 µL of RNase-free water. DNA was depleted by adding 2 µL of TURBO DNase and 10 µL of 10X Turbo DNase Buffer (ThermoFisher cat. #AM1907) and incubated at 37 °C for 15 minutes before being processed using MEGAclear™ Transcription Clean-Up Kit (ThermoFisher cat. #AM1908) following suggested protocols. Samples were mixed with Binding solution and 100% EtOH applied to filter cartridges and centrifuged. Filter cartridges were washed twice with Wash Solution and centrifuged to remove any traces of Wash Solution. DNA-depleted RNA was eluted in 50 µL of RNAse-free water. 500 ng of purified RNA was added to a master mix containing 10 pmol of dNTPs, 20 pmol of reverse primer specific to genomic MS2 RNA, and molecular grade water to reach 14.5 µL. The mixture was incubated at 95 °C for 5 minutes and then immediately placed on ice. To each sample, 20U of SUPERase·In™ RNase Inhibitor (ThermoFisher cat. #AM2696) along with 100U of Maxima H Minus Reverse Transcriptase and 5X RT Buffer (ThermoFisher cat. #EP0752) was added and samples were incubated in the thermocycler using the following program: [65 °C for 30 min, 85 °C for 5 minutes]. Samples were treated with 10U of RNase H (ThermoFisher cat. #AM2293) and incubated at 37 °C for 20 minutes. Samples were processed via DNA Clean & Concentrator-5 kit (Zymo Research cat. #D4003) as per the manufacturer’s instructions indicated for cDNA samples. Purified cDNA underwent qPCR using 300 nM of forward and reverse primers specific to genomic MS2 RNA, 250 nM of probe containing a 5’-6-FAM modification, a ZEN internal quencher modification, and a 3’-Iowa Black fluorescent quencher modification, and iTaq Universal Probes Supermix (BioRad cat. #1725130). Samples were incubated in the QuantStudio3 Real-Time PCR analyzer using the following program: [Step 1. 95 °C for 2 minutes; Step 2. 95 °C for 15 seconds, Step 3. 60 °C for 60 seconds, Step 4. Return to Step 2 for 39X]. A standard curve was used to quantify MS2 gRNA levels. Standards were generated by serial diluting known concentrations of a gBlock containing the target sequence for MS2 gRNA. Analysis of qPCR data was performed using QuantStudio Design and Analysis (DA) Software. See **Extended Data Table 4** for sequences and modifications of primers and probes used.

### One-Step Growth Curves

Constructs containing *gfp* or *ISG20*_*WT*_ were expressed and passaged as described in the first paragraph of the Methods section. Mid-log cells (OD_600_ ~0.7) were dosed with phage in 10 mL culture volumes at an MOI of 0.1. Media only controls were made by inoculating the same amount of phage to 10 mL of fresh media to serve as phage input controls. Phage mixtures were incubated shaking at 37 °C for 15 minutes before centrifugation (4000g, 10 min) at room temperature. Supernatants from *gfp* and *ISG20*_*WT*_ were collected for titering unbound phages, and media-only controls were used to measure input phage counts. The pellet of *gfp* and *ISG20*_*WT*_ cells was resuspended in 10 mL of fresh media and immediately serial diluted (tenfold dilutions) into three 25ml flasks containing growth media. At indicated timepoints, 100 µL was taken from each flask, mixed with 100 µL of our indicator strain, *gfp-Turbo*, and 3 mL of LB top agar, and poured onto LB agar plates containing kanamycin (50 μg mL^−1^) and IPTG (0.5 mM). After plating, samples were dried by flame for ~5 minutes before incubation at 37 °C overnight. Plaques were quantitated for each dilution at each timepoint to calculate the PFU/mL over one burst cycle of phage. The number of phage-bound cells was calculated by subtracting the unbound phages from the input phage. Emergent PFU at each timepoint was normalized to the number of phage-bound cells.

### Protein expression and purification

Recombinant ISG20_WT_ and ISG20_DEAD_ was expressed and purified as previously described, with minor modifications^34^. Briefly, the coding sequence of ISG20 was codon-optimized for *E. coli* per **Extended Data Table 5** and cloned with a C-terminal hexa-histidine tag into our modified pDEST14 vector (ThermoFisher cat. #11801016). ISG20_WT_ and ISG20_DEAD_ constructs were chemically transformed into *E. coli* BL21-CodonPlus (DE3)-RIPL (Agilent) before single colonies were grown overnight in LB media containing 200 ug/mL of carbenicillin (LB-carb). The next day, 2% outgrowth cultures were made in two, 1 L flasks of fresh LB-carb media and grown to an OD_600_ value of ~0.6-0.7, before one was induced with 50 µM IPTG and the second induced with 500 µM of IPTG. Induced cultures were incubated at 16 °C overnight, shaking at 200rpm. The next day, pellets were collected after centrifugation (4500g, 12 min) at 4 °C and stored at −20 °C. Pellets were resuspended in buffer containing 50 mM Tris-HCl pH7, 50 mM of NaCl, 1 µM Betamercaptoethanol, 10% glycerol, and 1 tablet/50 mL Pierce protease inhibitor tablets (ThermoFisher cat. #A32963). Resuspended pellets from each flask were combined, sonicated (Qsoncia cat. #Q125, 60% amplitude, 8 pulses of 15s ON / 59s OFF), and centrifuged (30000g, 30 min) at 4 °C. Cleared lysates were supplemented with 5 mM imidazole and 500 mM NaCl. 1 mL of equilibrated 50% HisPur Ni-NTA bead suspension (ThermoFisher cat. #88221) was added to the lysate and incubated at 4 °C for at least 1 hour while rocking. Recombinant ISG20 was purified over a column. The column was washed in succession with the following buffers: Wash 1 (50 mM Tris-HCl pH 7, 1mM Betamercaptoethanol, 500 mM NaCl, 25 mM imidazole, 2 mM ATP, 10 mM MgCl_2_), Wash 2 (50 mM Tris-HCl pH 7, 1 mM Betamercaptoethanol, 1M NaCl, 25 mM imidazole), Wash 3 (50 mM Tris-HCl pH 7, 1 mM Betamercaptoethanol, 100 mM NaCl, 25 mM imidazole), and Wash 4 (50 mM Tris-HCl pH 7, 1 mM Betamercaptoethanol, 500 mM NaCl, 25 mM imidazole). Proteins were eluted using Elution Buffer containing 50 mM Tris-HCl pH 7, 1 mM Betamercaptoethanol, 250 mM NaCl, and 250 mM imidazole. Elutions were concentrated on a 10 kD MWCO Amicon filter before undergoing size exclusion chromatography using SEC Running Buffer (50 mM Tris-HCl pH 7, 1 mM Betamercaptoethanol, and 250 mM NaCl). Eluted proteins were concentrated on a 10 kD MWCO Amicon filter before being flash frozen in liquid nitrogen and stored at −80 °C.

The coding sequence of EcTon followed by a C-terminal Twin-Strep-tag was cloned into our pET15 vector (**Extended Data Table 5**). EcTON_WT_ and EcTON_DEAD_ constructs were chemically transformed into One Shot™ BL21(DE3)pLysS *E. coli* (ThermoFisher cat. #C606003) before single colonies were grown overnight in LB-carb media. The next day, 3% outgrowth cultures were made in 1L of fresh LB-carb media and grown to mid log phase (OD_600_ ~ 0.5-0.6) before being induced with 500 uM of IPTG for 2.5 hours. Pellets were collected and stored as described above. Pellets were resuspended in 40 mL of Resuspension Buffer (100 mM Tris-HCl, pH 8.0, 300 mM NaCl, 1 mM EDTA, 1 mM DTT, 1 tablet/50 mL Pierce Protease Inhibitor tablets) before adding 0.1 mg/mL of lysozyme and incubating on ice for 10 minutes. Samples were sonicated as for ISG20 and passed through a 18g needed 2-3 times to sheer genomic DNA and decrease sample viscosity. Samples were centrifuged (30000g, 30 min) at 4 °C and clarified lysates were passed through a 0.45 um filter before loading onto purification columns. EcTON_WT_ and EcTON_DEAD_ proteins were purified on Strep-Tactin®XT 4Flow gravity flow columns (IBA cat #2-5013-001) following modified manufacturer’s instructions. Briefly, the column was washed with 5 column volumes (CV) of Wash Buffer (100 mM Tris-HCl, pH 8.0, 300 mM NaCl, 1 mM EDTA, 1 mM DTT), 2 CV of ATP Wash Buffer (100 mM Tris-HCl, pH 8.0, 300 mM NaCl, 1 mM EDTA, 1 mM DTT, 2 mM ATP, 10 mM MgCl_2_) and another 2 CV of Wash Buffer. Proteins were eluted in 5 CV of 1X BXT Buffer (IBA cat# 2-1042-025). Elution was concentrated on a 10 kD MWCO Amicon filter before undergoing size exclusion chromatography using high salt SEC Running Buffer containing 50 mM Tris-HCl pH 7, 1 mM Betamercaptoethanol, and 500 mM NaCl. Eluted proteins were concentrated on a 10 kD MWCO Amicon filter before being flash frozen in liquid nitrogen and stored at −80 °C.

### Nuclease Assays

Nuclease assays were performed by mixing recombinant ISG20_WT_, EcTON_WT,_ or matching catalytically dead mutants at concentrations 500 nM, 100 nM, 20 nM, and 4 nM, in a 4X Nuclease Buffer (200 mM Tris-HCl pH7, 10 mM MnCl_2_, 4 mM Betamercaptoethanol, 1.6 mM DTT, 0.4% Triton X-100, 0.2 mg/mL Bovine Serum Albumin, and 40% glycerol). Synthesized RNA substrates containing a 5’ 6-FAM fluorescent label were melted and annealed in a thermocycler using the following program: [95 °C for 5 minutes, 25 °C ramp 0.1C/sec, 25 °C for 20 minutes, 12 °C hold] before being added to the nuclease reaction at final concentration of 500 nM. The reaction was incubated at 37 °C for 15 minutes before 8 µL of the reaction was transferred to 8 µL of Gel Loading Buffer II (ThermoFisher cat. #AM8546G) to stop the reaction. The collected samples were incubated at 98 °C for 5 minutes then immediately placed on ice before being resolved on a 15% TBE-urea polyacrylamide gel for 80 minutes at 200V. Gels were stained with Sybr Gold and imaged on the ChemiDoc imaging system. See **Extended Data Table 4** for sequences and modifications of RNA substrates used.

### 3’ UTR Sequencing

We sequenced MS2 3’ UTRs by adapting protocols for 3’ Rapid Amplification of cDNA Ends (3’ RACE). Nuclease assays were performed, as previously described above, using 2.5 µM of recombinant ISG20_WT_ protein and 2.5 µM of MS2-UTR RNA substrate incubated for 15 minutes at 37 °C and resolved on a 15% TBE-urea polyacrylamide gel for 80 minutes at 200V. Gels were stained with Sybr Gold and imaged on the ChemiDoc imaging system. RNA gel bands were excised and processed via ZR small-RNA PAGE recovery kit (Zymo Research cat. #R1070) following manufacturer’s instruction. RNA adapter containing a 5’-phosphate and 3’-inverted thymidine was ligated to purified RNA using Ambion T4 RNA ligase kits (ThermoFisher cat. #AM2141) following the manufacturer’s instruction. Ligase reactions were incubated at 4 °C overnight. Molecular grade water was added to samples to a total volume of 50 uL and processed using the Zymo RNA Clean and Concentrator −5 kit (Zymo Research cat. #R1013) following the manufacturer’s instruction. cDNA was synthesized as described above. Reverse primers specific to the 3’ region of the RNA adapter were used for reverse transcription. Following reverse transcription, samples were treated with RNase H and processed via ZymoClean DNA Clean and Concentrator kit-5 as described above. Purified cDNA was PCR amplified using 200 nM of forward primer specific to 3’-UTR of MS2, 200 nM of RNA adapter-specific reverse primer, cDNA, Q5® High-Fidelity Master Mix (NEB cat. #M0492S), and molecular grade water. Samples were incubated in the thermocycler using the following program: [Step 1. 98 °C for 30 seconds; Step 2. 98 °C for 10 seconds, Step 3. 67 °C for 30 seconds, Step 4. 72 °C for 10 seconds, Step 5. Return to Step 2 for 39X, Step 6. 72 °C for 2 min, Step 7. 4 °C Hold.]. Samples were processed via DNA Clean & Concentrator-5 kit (Zymo Research cat. #D4003) as per the manufacturer’s instructions indicated for cDNA samples before being sent out to Plasmidsaurus for Premium PCR sequencing. Oligonucletide sequences and modifications are listed in **Extended Data Table 4**.

To quantify 3’ UTR sequences, fastq reads were mapped to the UTR-adapter hybrid using the Geneious Prime sequence analysis suite (v2022.2.2), assuming a wild-type sequence. Mapped reads were then extracted and considered further if they contained the final nine bases of the MS2-trunc cDNA sequence (CTAGTTACC) and the first nine of ligated RNA adapter (TCACACTGA). This approach ensured that only appropriately ligated UTR sequences were considered. To quantify which of the remaining five ribonucleotides (ACCCA) remained associated with the 3’ UTR, if any, sequencing reads were tallied according to their intervening sequence. The percentage of reads with each possible overhang are depicted in **Figure 1h**.

### Homology Searches

The human ISG20 protein was used as an initial query in all searches (NCBI ID: NP_002192.2). To search NCBI’s nr and nt databases, blastp and tblastn were used, respectively. In both cases, searches were performed in the summer of 2022. The top 50 hits under 500 amino acids, assessed by e-value, were examined manually to ensure true bacterial origin (for instance, by identifying multiple homologous sequences in high-quality bacterial genome assemblies). To query NCBI’s conserved domain database, ISG20 was identified within the conserved domain family ‘cd06127’, which corresponds to DEDDh family exonucleases (ISG20 falls within the sub-clade cd06149). The .cn4 phylogeny for cd06127 was downloaded from NCBI and the cladogram analyzed backwards from cd06149, looking for representative sequences of bacterial origin as the evolutionary history of this clade was traced towards its ancestor. Clades cd06144, cd06143, and cd06137 were eukaryote-specific. Bacterial sequences were first encountered in the clade cd06126. These sequences were downloaded, examined manually as before to ensure bacterial origin, and combined with those retrieved via blast for further analysis.

Candidate homologs were aligned using MUSCLE with default parameters and the DEDDh domain extracted by manual examination in the Geneious Prime sequence analysis suite (v2022.2.2) ^47^. A preliminary phylogenetic tree was then generated from this alignment using PhyML^48^. Sequences for follow-up were then selected to cover the phylogenetic diversity present in the initial hit list. Preference was given for sequences from gammaproteobacteria and those nearby predicted phage defense genes, predicted phage genes, or other functions associated with horizontal gene transfer. This enriched set was re-aligned, new phylogenetic trees created, and a representative set of 27 synthesized in pZE21_TetR by GenScript for functional testing (**Extended Data Tables 2 and 4**). To identify structural homologs of EcTon, AlphaFold3 was used to model its tertiary structure, which produced a predicted structure with high confidence (pTM = 0.79). This structure was used as a query in a DALI search of the PDB90 database (search data: April 22, 2026) ^49^. **Extended Data Table 3** reports the top ten high-confidence hits from this search (i.e. those with an RMSD score better than 5).

### Phylogenetic Analysis

The 27 bacterial homologs were aligned by muscle and the DEDDh domain extracted by manual examination in the Geneious Prime sequence analysis suite (v2022.2.2). This domain was re-aligned by MUSCLE and a phylogenetic tree created using PhyML with the LG substitution model with 100 bootstraps to create the tree depicted in **Figure 3a**^48^.

### Cell lines and cell culture

Mammalian cell lines were maintained at 37 °C in 5% CO_2_ atmosphere. U2OS cells were obtained from ATCC (HTB-896) and cultured in DMEM (Sigma) supplemented with 10% FBS. BSC-40 cells were cultured in MEM (Sigma) supplemented with 5% FBS. All media additionally contained 1% non-essential amino acids, 1% L-glutamine, and 1% antibiotic/antimycotic (Gibco).

### Mammalian Viruses

VEEV-GFP^41^, ONNV-GFP^41^, VSV-GFP^42^, and VACV-FL-GFP^42^ stock preparations and titration by fluorescent foci/plaque assay on BSC40 cells were performed as previously described^40–42^. Viral inocula were incubated with cells for 1 hour in serum free media before the addition of complete media for the remainder of the infection as previously described^41,42^.

### General transfection and expression in mammalian cells

Bacterial TONDOs and human ISG20 were codon-optimized for mammalian expression before synthesis and cloning into pcDNA3.1 (performed by GenScript Biotech). The pCDNA3.1 expression plasmid contained a C-terminal FLAG epitope tag. Unless otherwise stated, 100,000 cells were transfected with 500 ng of expression vectors (**Extended Data Table 5**) and expressed for 24 hours. For U2OS cell expression, pcDNA3.1 constructs were mixed with 100 μL of OptiMEM (Gibco) media and 1.5 μL Lipofectamine 2000 reagent (Thermo Fisher). This mixture was incubated at room temperature for 20 minutes, the media on the cells was changed to 400 μL Opti-MEM, and then the transfection mix was added dropwise to the cells and incubated for 6 hours. Following transfection, cell media was changed to complete DMEM and expression progressed for 24 hours prior to viral challenge or immunoblotting.

### Immunoblotting

Protein extracts were generated via lysis of U2OS cells in RIPA buffer (Thermo Fisher, cat# 89901) and diluted in 5X SDS-PAGE loading buffer then boiled at 95 °C for 10 minutes. Samples were subjected to SDS-PAGE electrophoresis at 100 V for approximately 1.5 hours. Separated proteins were transferred to nitrocellulose membranes in 1X transfer buffer (BioRad cat #1704271) at 1300 mA at 25 °C. Membranes were blocked in EveryBlot Blocking Buffer (BioRad, cat #12010020) for 1 hour at 25 °C. Membranes were blotted with primary antibody overnight at 4 °C (Mouse-anti-Actin: Sigma Cat# A1978, Rabbit-anti-Flag: Thermo cat #740001), with actin serving as a loading control unless otherwise stated. The following morning, three, 5 min washes with PBS-T (PBS, 0.1% Tween, BioRad cat #1610780), membranes were incubated in secondary antibody (BioRad cat #12004159 for anti-mouse, or 12005870 for anti-rabbit) for 1 hour followed by three, 5 min washes in PBS-T and one 5 min PBS wash. Membranes were then imaged via BioRad Imager.

### Cell viability assay

The toxicity of the pcDNA3.1 ISG20 constructs with was measured using a CyQUANT LDH Cytotoxicity assay (Invitrogen). Briefly, 100,000 U2OS cells per well were plated into 24-well plates and transfected the next day. Following 24 hours of expression, 25 μL of supernatant from transfected cells was transferred to a fresh 96-well plate. Then, 25 μL of Lactose Dehydrogenase (LDH) reaction mixtures was added to each well (Thermo Fisher CyQUANT LDH cytotoxicity assay, cat. #C20300). The plates were mixed gently by tapping ~5 times and incubated in the dark for 30 min. Then, 25 μL of Stop Solution was added (included in CyQUANT kit). The absorbance of each plate was read at both 490 nm and 680 nm within 15 minutes of adding Stop Solution. Data plotted indicates the 680 nm value (background signal of instrument) subtracted by the 490 nm value, per manufacturer’s instructions. Controls included: DMEM “media-only” negative control, and a “lysed cells” positive control of U2OS cells. Positive control cells were lysed per manufacturer’s instructions one hour before collecting supernatant media.

### Statistical Analyses

Viral infection and cell viability graphs are presented as mean values ± SD with individual data points shown. At least three independent biological replicates were conducted for all quantitative experiments (those with statistics provided). All statistical analyses were performed with Prism software v10.0.2 (GraphPad) or Excel. Statistical tests used are indicated in respective figure legends.

## Acknowledgements

We thank John Schoggins for early conversations on ISG diversity and function. We also thank Cassie Alvorado and Amanda Dobbins for helping purify ISG20, Senen Mendoza for qRT-PCR protocols, and Amitabh Ranjan for advice on UTR sequencing; from the labs of Xinyun Cao, David Hendrixson, Michael Laub, and Nick Conrad, respectively.

## Author Contributions

Conceptualization, K.J.F.; methodology, C.R.S., A.E-G., A.E., and K.J.F.; investigation, C.R.S., A.E-G., A.E., A.E.M., and K.J.F.; funding acquisition, N.M.A., D.B.G., and K.J.F.; project administration, N.M.A., D.B.G., and K.J.F.; supervision, D.B.G. and K.J.F.; writing—original draft, C.R.S. and K.J.F.; writing—review and editing, C.R.S., A.E-G., A.E., D.B.G., and K.J.F.

---

**Extended Data Fig. 1.**
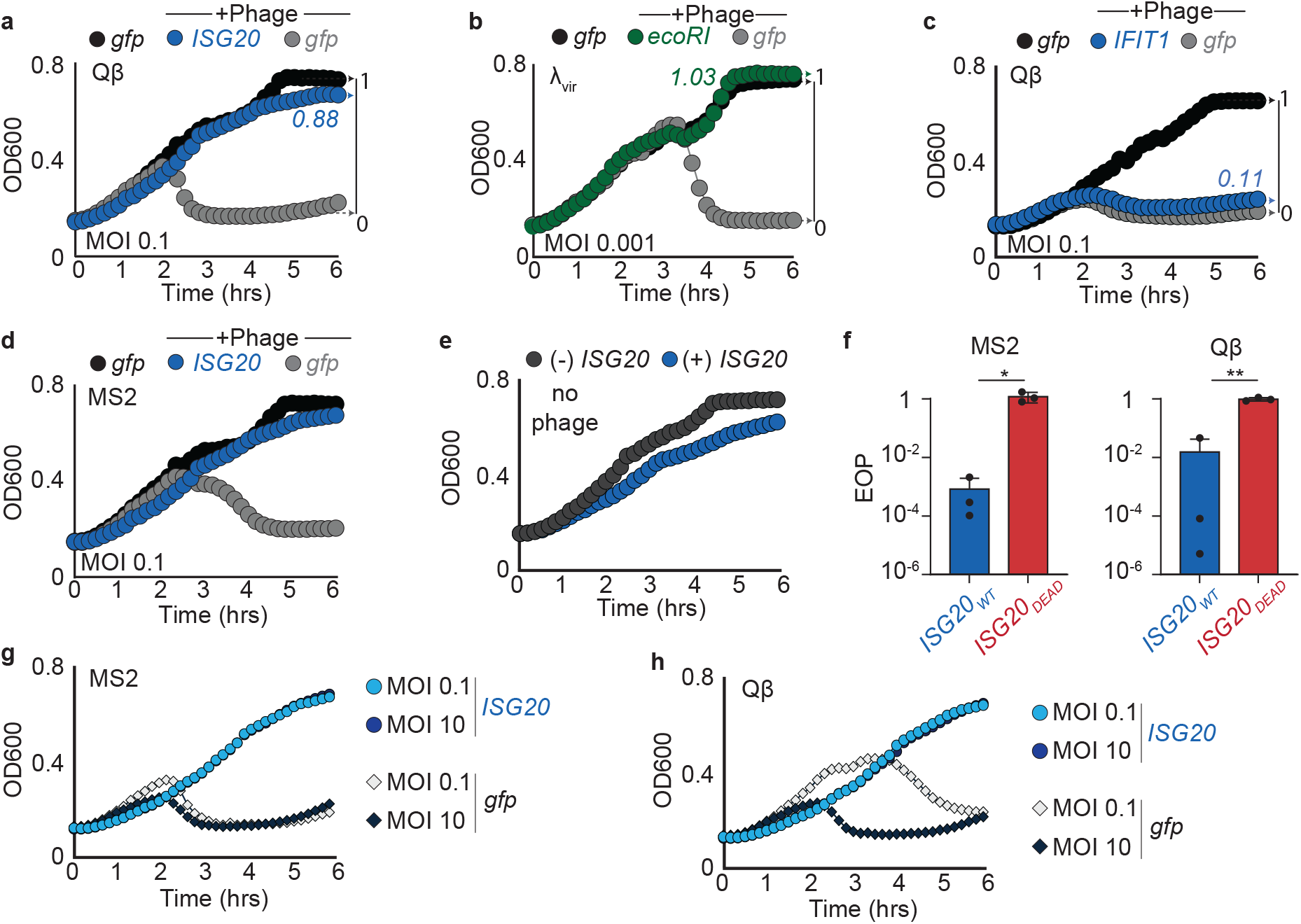
Phage defense phenotypes for various immunity genes. **a**, Growth curves of *E. coli* expressing *gfp* or *ISG20* in the presence or absence of Qβ as indicated in the legend at top. The number line at right provides scale for the protection score assigned to *ISG20* from this growth curve, in colored numerals and shown in **Figure 1b. b**, Growth curves of *E. coli* expressing *gfp* or *ecoRI* in the presence or absence of λ_vir_ as indicated in the legend at top. **c**, Growth curves of *E. coli* expressing *gfp* or *IFIT1* in the presence or absence of Qβ as indicated in the legend at top. Protection scores in panels **b** and **c** as in panel **a. d**, Growth curves of *E. coli* expressing *gfp* or *ISG20* in the presence or absence of MS2 as indicated in the legend at top. **e**, Growth curves of *E. coli* expressing *ISG20* compared to controls for which expression is not induced. **f**, Efficiency of plaquing (EOP) for MS2 (left) or Qβ (right) on *E. coli* expressing the indicated genes. *ISG20*_*WT*_ and *ISG20*_*DEAD*_ EOP values are calculated relative to matched GFP controls, which was set to 1. The data represent averages of 3 biological replicates ± s.e.m. Statistical significance determined by Welch’s T-test; *p≤0.05; **p≤0.005. **g**, Growth curves of *E. coli* expressing *ISG20* or *gfp* in the presence of MS2 at MOI of 0.1 or 10. **h**, Growth curves of *E. coli* expressing *ISG20* or *gfp* in the presence of Qβ at MOI of 0.1 or 10. OD600 signifies optical density measurements, determined via absorbance of 600nm light. Growth curves depict averages of three technical replicates and are representative of three biological replicates. Growth curve error bars plot st. dev but are typically obscured by growth curve data points.

**Extended Data Fig. 2.**
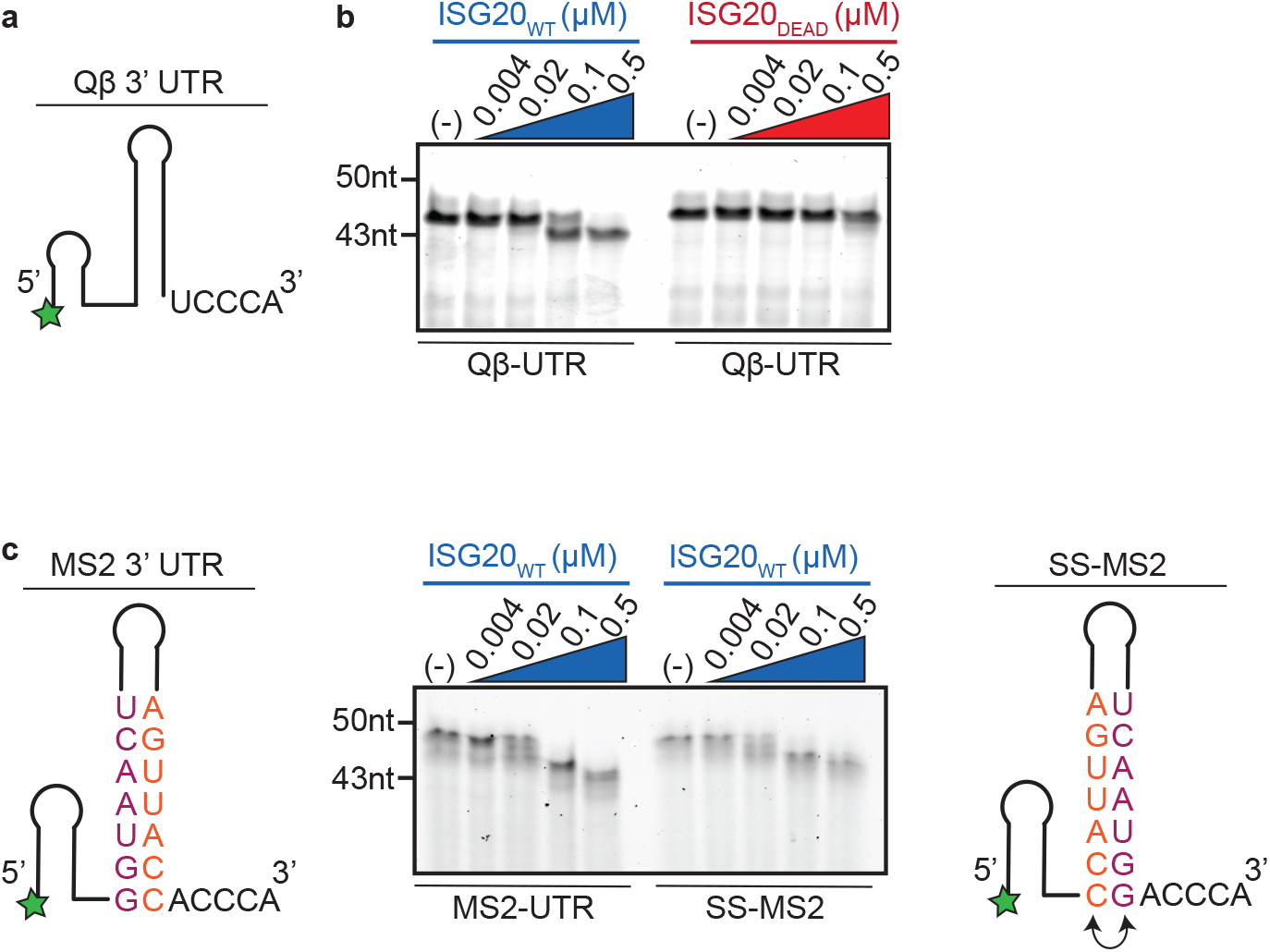
ISG20 degrades MS2 and Qβ 3’ UTRs. **a**, Diagram of the terminal region of Qβ’s 3’ UTR, synthesized with a 5’ 6-FAM fluorophore (green star). **b**, Degradation of the Qβ-UTR (0.5µM) after 15-minute incubations with either ISG20_WT_ or ISG20_DEAD_, at the indicated protein concentrations. **c**, Degradation products from the MS2-UTR and SS-MS2 substrates (0.5µM) incubated with ISG20_WT_, as in panel **b**. Schematics depict each substrate to the left and right of the gel. RNA gels are representative of three biological replicates.

**Extended Data Fig. 3.**
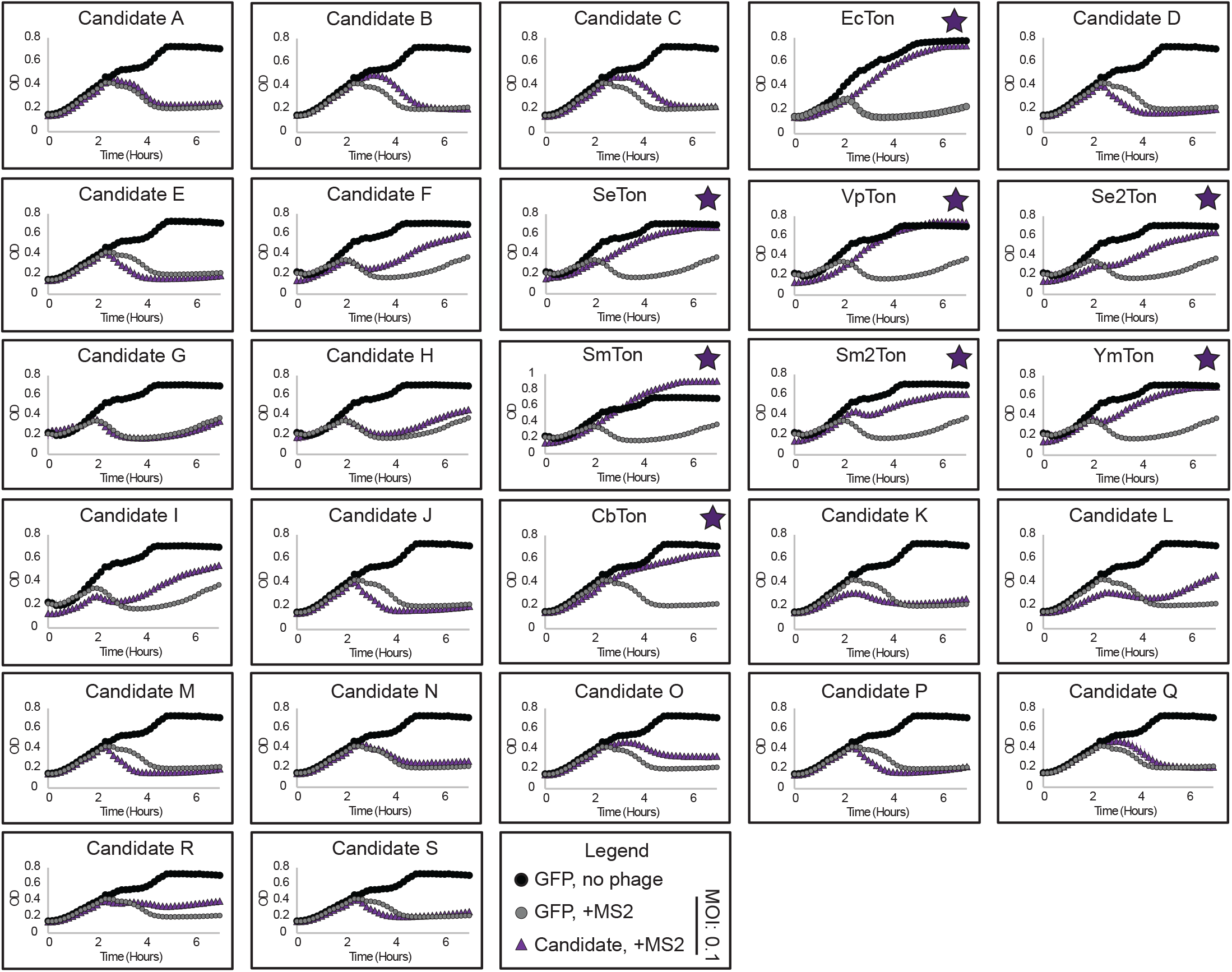
Some bacterial homologs of ISG20 protect *E. coli* from MS2 infection. Growth curves of candidate antiphage exonuclease genes (purple) in the presence of MS2 (MOI 0.1). Curves are compared to *gfp*-expressing cells in the presence (gray) or absence (black) of phage. Genes with antiphage defense phenotypes were named TONDOs and are indicated by purple stars. Legend right of bottom row. OD signifies optical density measurements, determined via absorbance of 600nm light. Growth curves depict averages of three technical replicates. TONDO curves are representative of three biological replicates. Error bars plot st. dev but are typically obscured by growth curve data points.

**Extended Data Fig. 4.**
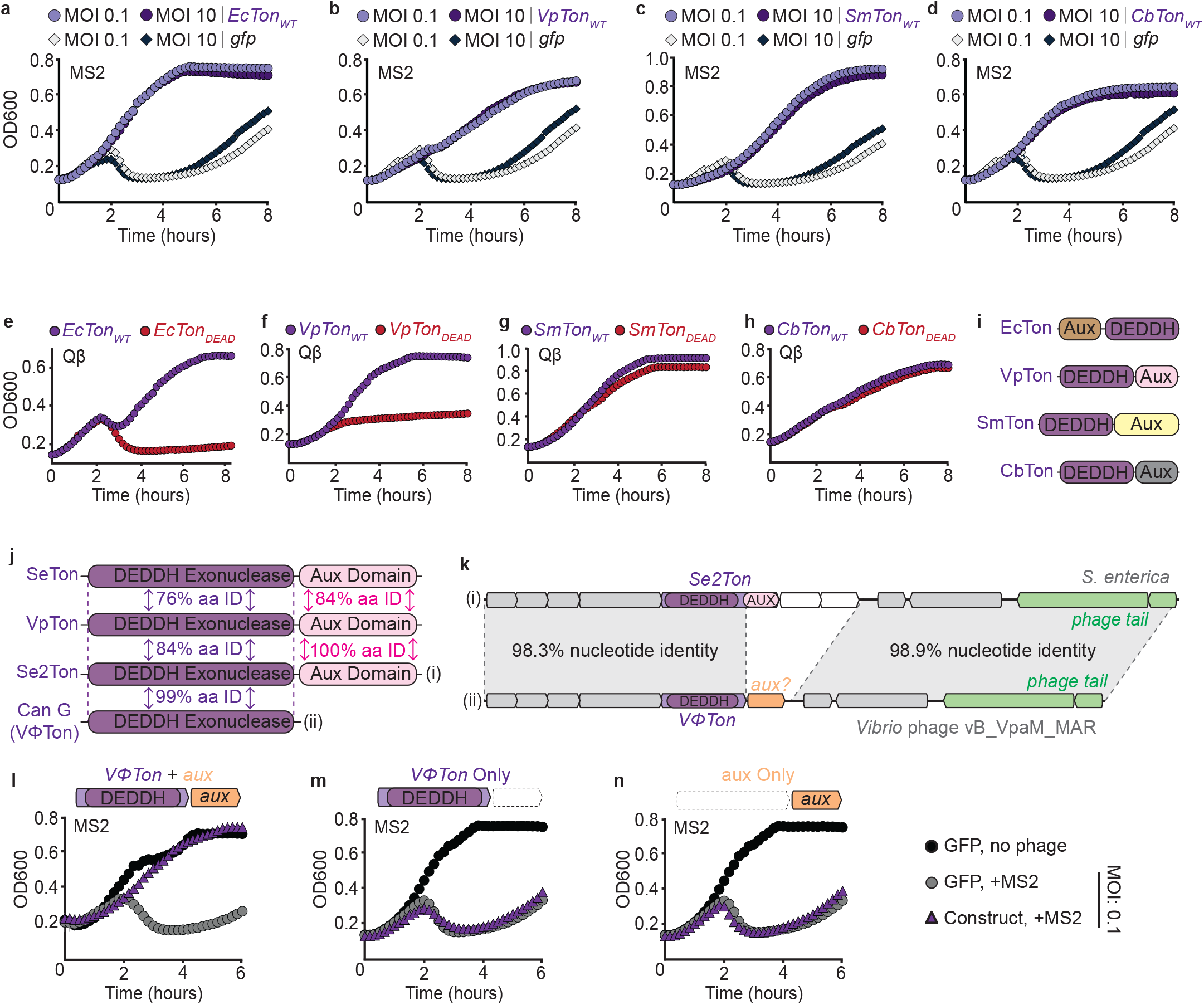
Genetic and phenotypic features of TONDOs. **a**, Growth curves of *E. coli* expressing *EcTon* or *gfp* in the presence of MS2 at MOI of 0.1 or 10. **b**, Growth curves of *E. coli* expressing *VpTon* or *gfp* in the presence of MS2 at MOI of 0.1 or 10. **c**, Growth curves of *E. coli* expressing *SmTon* or *gfp* in the presence of MS2 at MOI of 0.1 or 10. **d**, Growth curves of *E. coli* expressing *CbTon* or *gfp* in the presence of MS2 at MOI of 0.1 or 10. **e-h**, Growth curves of *E. coli* expressing the indicated TONDO (purple) or its catalytically inactive mutant (red), in the presence of Qβ at an initial MOI of 0.1. **e**, Qβ defense by *EcTon*_*WT*_ and *EcTon*_*DEAD*_. **f**, Qβ defense by *VpTon*_*WT*_ and *VpTon*_*DEAD*_. **g**, Qβ defense by *SmTon*_*WT*_ and *SmTon*_*DEAD*_. **h**, Qβ defense by *CbTon*_*WT*_ and *CbTon*_*DEAD*_. **i**, domain organization of select TONDO genes. Auxiliary (aux) domains have no detectable homology so are colored differently. **j**, Pairwise amino acid identities of related TONDOs and related candidates. Candidate G (VΦTon) lacks the auxiliary domain present in its close relatives. **k**, *Se2Ton* and *VΦTon* are encoded by highly related predicted prophages but differ in the sequence immediately downstream of the TONDO gene. **l**,**m**,**n**, Growth curves of the indicated genes (purple) in the presence of MS2 (MOI 0.1). Curves are compared to *gfp*-expressing cells in the presence (gray) or absence (black) of phage, per the legend at right. OD600 signifies optical density measurements, determined via absorbance of 600nm light. Growth curves depict averages of three technical replicates and are representative of three biological replicates. Error bars plot st. dev but are typically obscured by growth curve data points.

**Extended Data Fig. 5.**
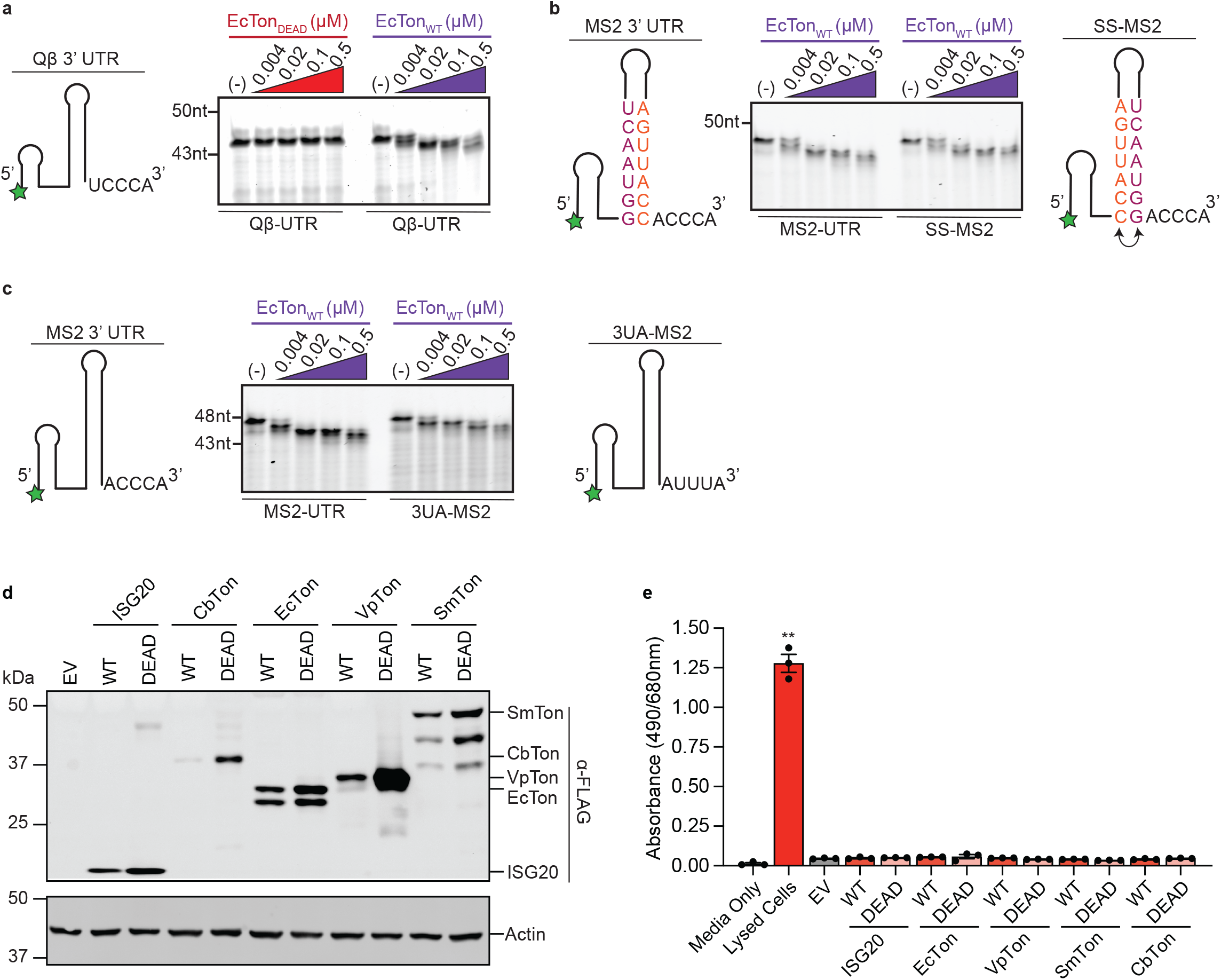
Enzymatic and cell biological features of TONDOs. **a**, Degradation of the Qβ-UTR (0.5µM) after 15-minute incubations with either EcTon_DEAD_ or EcTon_WT_ at the indicated protein concentrations. **b**, Degradation products from the MS2-UTR and SS-MS2 substrates (0.5µM) incubated with EcTon_WT_. **c**, Degradation products from the MS2-UTR and 3UA-MS2 (0.5µM) substrates incubated with EcTon_WT_. Schematics adjacent to gels depict the UTR substrates used. **d**, Representative immunoblot of U2OS cells expressing the indicated ISG20 or TONDO construct with a C-terminal FLAG-tag. Bands smaller than full-length TONDOs may represent partially proteolyzed products. **e**, Lactose Dehydrogenase cytoxicity assay conducted on U2OS cell supernatant following 48 hours of expressing the indicated ISG20 or TONDO construct. The data represent means ± st. dev. (n=3 biological replicates). Statistical significance was determined by one-way ANOVA and Dunnett’s post-test, compared against media-only controls. Two asterisks (**) denote p≤0.005. Comparisons without asterisks are not significant.

**Extended Data Fig. 6.**
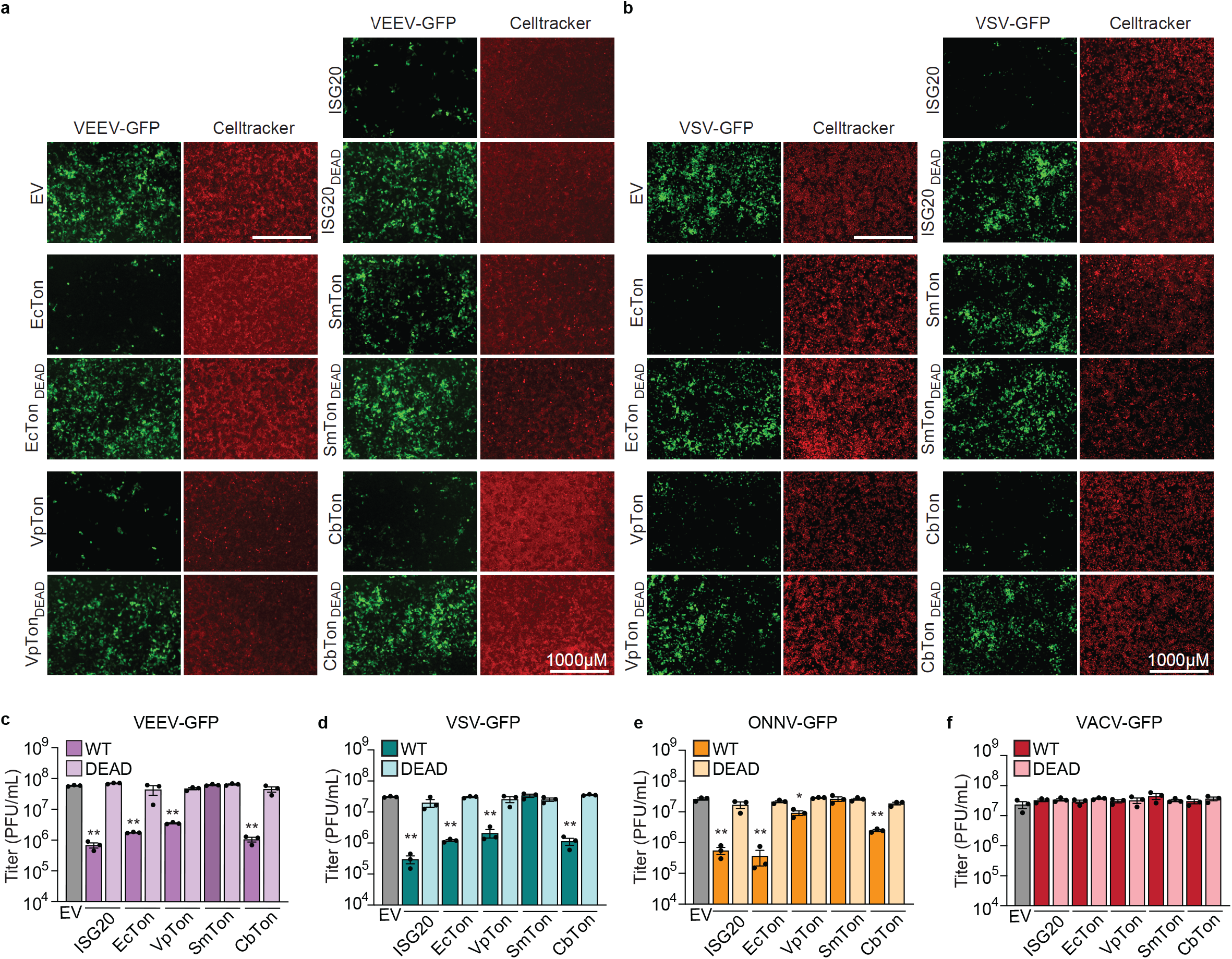
Impact of TONDO expression on human virus infections. **a**, Representative images of TONDO-expressing U2OS cells 24 hours post-infection (hpi) with VEEV-GFP (MOI 1). GFP signal (green) marks VEEV infection, Celltracker stain (red) marks cell area. Scale bar, 1000µm. The images are representative of three independent replicates, quantified in **Figure 4b. b**, Representative images of TONDO-expressing U2OS cells 24 hpi with VSV-GFP (MOI 1). GFP signal (green) marks VSV infection, Celltracker stain (red) marks cell area. Scale bar, 1000µm. The images are representative of three independent replicates, quantified in **Figure 4c. c**, Viral titers of supernatants collected from VEEV-GFP infected U2OS cells 24 hpi. **d**, Viral titers of supernatants collected from VSV-GFP infected U2OS cells 24 hpi. **e**, Viral titers of supernatants collected from ONNV-GFP infected U2OS cells 24 hpi. **f**, Viral titers of supernatants collected from VACV-GFP infected U2OS cells 24 hpi. Data represent means ± st. dev (n=3 biological replicates). Statistical significance determined by one-way ANOVA and Dunnett’s post-test, compared to EV controls; *p≤0.05, **p≤0.005. Comparisons without asterisks are not statistically significant.

